# A family of lethal exotoxins defined by cell entry via the Attractin receptor

**DOI:** 10.1101/2025.10.08.681221

**Authors:** Raghuvir Viswanatha, Donghoon Lee, William R. Robins, Enzo Mameli, Yanhui Hu, Ah-Ram Kim, Yousuf Hashmi, Hiroshi Nishida, Gyan Prakash, Matthew Butnaru, Sterling Churchman, Stephanie E. Mohr, John J. Mekalanos, Norbert Perrimon

**Author notes:** Corresponding authors: Norbert Perrimon, John Mekalanos, and Raghuvir Viswanatha. These authors contributed equally to this work (Raghuvir Viswanatha, Donghoon Lee, William R. Robins).

## Abstract

Although bacterial genomes encode numerous potential toxins, it is unclear how evolution drives the specificity of these important virulence factors. Using an insect CRISPR screen, we identified the transmembrane protein Attractin (ATRN) as the receptor for Nigritoxin (Ntx), a Vibrio toxin that causes seasonal shrimp pandemics. We found that Ntx’s effector “warhead” inhibits translation via a previously uncharacterized mechanism. Moreover, we show that two related toxins require ATRN for entry but possess unrelated effector domains. One has a Rho-GTPase AMPylation function and the other an actin-targeting/proteolysis function. Our findings reveal the mechanism of Ntx entry and toxicity and show that the ATRN-targeting domain can deliver disparate effector domains, strongly indicating that this class of exotoxins can evolve as modular proteins using a common entry domain.

---

Bacteria secrete protein toxins that target eukaryotic cells, causing modulation of cell functions, death and ultimately disease (*1, 2*). Despite the abundance of toxin-like genes found across bacterial genomes, current mechanistic insight comes primarily from a handful of well-studied archetypes such as cholera, diphtheria, botulinum, and anthrax. Toxins that contribute to diseases of arthropods and crustaceans are far less studied and usually investigated following major losses with economic impact. *Vibrio* species in particular are significant agents in aquaculture and food poisoning (*3, 4*), with lethal shrimp outbreaks alone costing billions of dollars annually (*5*). The genomes of many *Vibrio* species implicated in invertebrate diseases commonly have genes in accessory islands and on plasmids that encode virulence factors including toxins. *Vibrio nigripulchritudo* has recently been identified across the Indo-Pacific region as a major seasonal pathogen of the *Litopenaeus* shrimp (*6, 7*). Its encoded toxin, Nigritoxin (Ntx), implicated in shrimp disease was found to be distinct from all previously studied toxins, though with some homology to an uncharacterized region of the anti-feeding phage toxin Afp18 (*8, 9*).

Ntx, a single-chain lethal toxin produced by hypervirulent *V. nigripulchritudo* strains, is the likely causative agent of Summer Syndrome, an Indo-Pacific shrimp pandemic characterized by necrosis of internal tissues, death of immune cells, and mortality within 1-3 days (*6, 7*). Initial characterization revealed that recombinant Ntx is lethal to crustaceans and insects but not mammalian cells (*9*). Structural studies showed that Ntx has three domains: a putative receptor-binding domain at the N-terminus, a middle domain putatively involved in endosome escape, and a unique protein fold in the C-terminus that induces apoptotic death (*9*) . Despite these insights, the receptor for Ntx and the molecular mechanisms underlying its cell entry and cytotoxicity remained elusive. Furthermore, its evolutionary history remains mysterious due to a lack of characterized orthologs.

## Identification of genes required for Ntx toxicity reveal the toxin receptor ATRN

Unbiased, genome-wide CRISPR screens provide a powerful approach to identify toxin receptors and mechanisms of action. We established a CRISPR screening platform in a *Drosophila melanogaster* cell line and recently expanded the technique to mosquito cell lines (*10-12*) . To select the optimal system for Ntx resistance screening, we measured the sensitivity of screening-compatible cell lines to Ntx. Sf9 from *Spodoptera frugiperda* moths were highly sensitive, as previously demonstrated (*9*), with an IC50 of 3 pM. We found that Sua-5B cells from *Anopheles* mosquitoes are also sensitive to Ntx, with an IC50 of 9 pM; whereas S2 cells are less sensitive, with an IC50 >10 nM (Fig. 1A; fig. S1A). We performed a genome-wide resistance screen by exposing Sua-5B cells expressing sgRNAs (∼100,000 sgRNA library targeting each gene with a modal number of 7 independent single-guide RNAs [sgRNAs]) to 3 nM Ntx for 2 weeks and harvested resistant cells. The screen revealed strong enrichment of sgRNAs targeting the gene encoding the mosquito ortholog of the mouse gene *Attractin* (Ag*ATRN, AGAP003506*) with 6/7 independent sgRNAs highly enriched in the resistant population (Fig. 1B; table S1), suggesting that loss of ATRN function confers resistance to Ntx. Other genes identified included orthologs of ubiquitination, glycosylation, protein folding, and membrane trafficking genes (Fig. 1C). Mouse ATRN is heavily glycosylated; thus, glycosylation genes are likely required for proper ATRN expression, recognition, or both (*13, 14*). The fly and mouse orthologs of one of these genes, the E3 ubiquitin ligase MGRN1, has been connected to ATRN (*15, 16*). In mice, *ATRN* and *MGRN1* function together in pigment-type switching, a process involving endolysosomal trafficking of ATRN from the limiting membrane of melanocytes to lysosomes (*15, 17*), suggesting that ubiquitylation genes identified in the screen are acting through MGRN1’s influence on ATRN trafficking. Consistent with this, knockout of the *MGRN1* ortholog in insect cells confers partial resistance to Ntx (fig. S1B), and overexpression of *MGRN1* in insect cells dramatically redistributes ATRN (fig. S1C). Thus the CRISPR screen identifies ATRN and genes that presumably act through ATRN to confer susceptibility to Ntx.

**Fig. 1.**
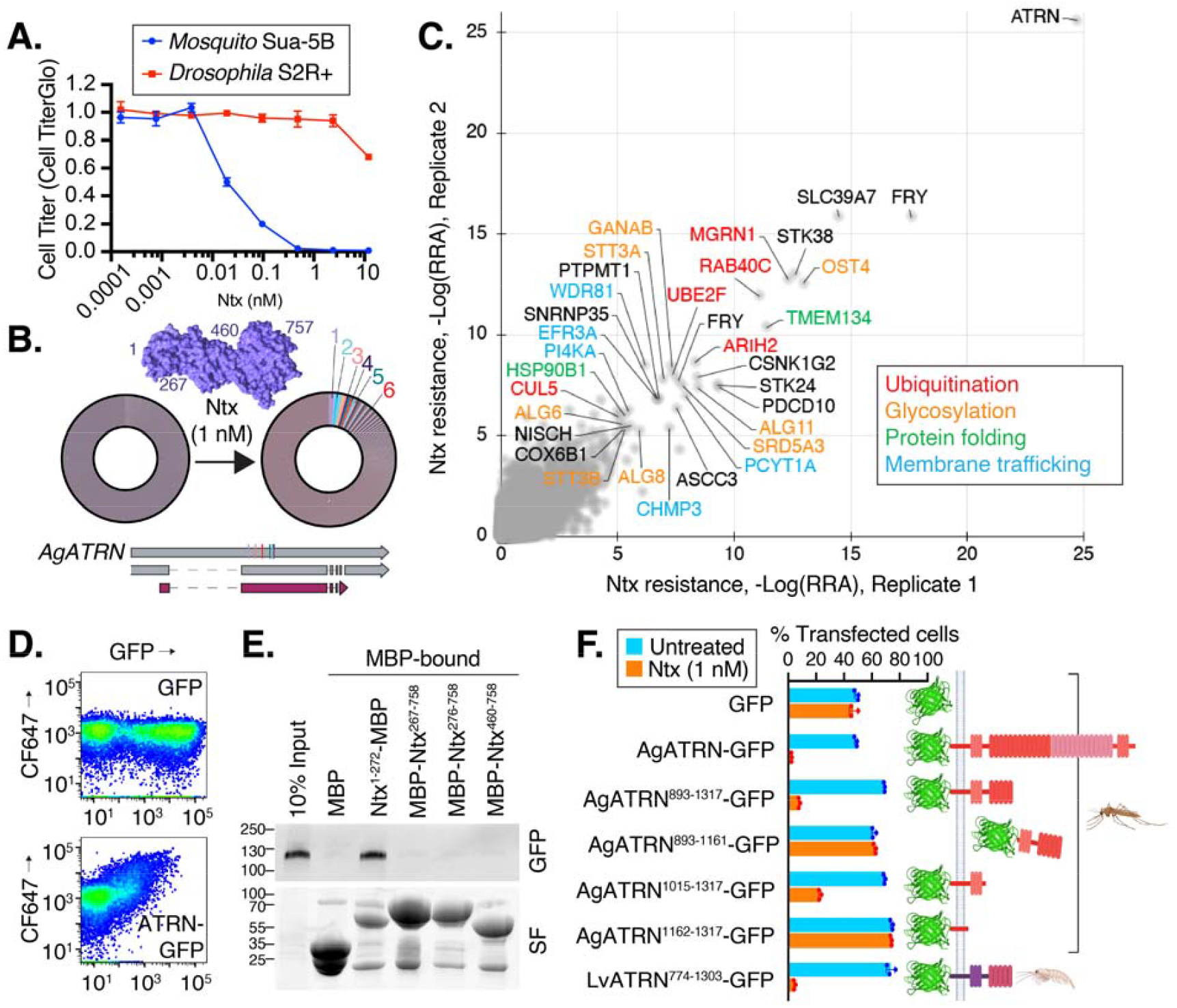
ATRN is the receptor for Nigritoxin (Ntx). **(A)** Cytotoxicity of Ntx in mosquito Sua-5B and *Drosophila* S2R+ cells. Viability was measured by CellTiter-Glo assay after 48 hours. Data are means ± SD (*n* = 3). **(B)** Donut charts showing the enrichment of sgRNAs targeting *ATRN* after Ntx selection. The left chart represents the initial plasmid library. The right chart shows the sgRNA population after selection, with 6/7 independent sgRNAs targeting *ATRN* (colored slices) highly enriched. Inset: Crystal structure of Ntx (PDB: 5M41). **(C)** Genome-wide CRISPR screen identifies ATRN as the top Ntx resistance hit as well as genes involved with ubiquitination, glycosylation, protein folding, and membrane trafficking. Scatter plot shows sgRNA enrichment from two replicate screens in Sua-5B cells treated with 3 nM Ntx. Hits are colored by functional category. **(D)** Ntx binds to cells expressing ATRN. Flow cytometry of cells expressing GFP or ATRN-GFP and incubated with fluorescently labeled Ntx (CF647-Ntx). **(E)** The N-terminal domain of Ntx binds directly to ATRN. MBP-fused Ntx fragments were used to pull down the GFP-tagged AgATRN. SF, stain-free gel, a total protein stain. **(F)** The ATRN stem domain is required for Ntx sensitivity. Moth Sf9 *ATRN* KO cells expressing the indicated ATRN constructs were treated with 1 nM Ntx. The percentage of GFP-positive cells with (blue bars) or without (orange bars) toxin treatment was quantified by flow cytometry. Data are means ± SD (*n* = 3).

To test if ATRN is the receptor for Ntx, we conducted cell-binding assays using cells expressing GFP-tagged *A. gambiae ATRN* (AgATRN-GFP). Expression of *AgATRN-GFP* led to a dose-dependent accumulation of fluorescently labeled Ntx (Fig. 1D; fig. S1D). Moreover, ATRN and Ntx co-immunoprecipitate in treated cell extracts (fig. S2E), and pulldown assays reveal a direct interaction between the N-terminal region of Ntx and ATRN (Fig. 1D). Together, these data suggest that the N-terminal domain of Ntx recognizes surface-exposed ATRN on insect cells.

## Identification of Ntx-binding region in the ATRN C-terminal stem domain

To pinpoint the region of ATRN responsible for Ntx binding, we utilized the more Ntx-sensitive Sf9 cell line with *ATRN* knocked out (fig. S2A) and reintroduced various GFP-tagged ATRN fragments. Flow cytometry-based cell competition assays revealed that cells expressing the ATRN stem domain (residues 1015–1317) regained sensitivity to Ntx, whereas cells expressing fragments lacking this domain did not (Fig. 1F). The equivalent domain of *L. vannemei* shrimp ATRN could also compensate for the loss of ATRN and confer sensitivity (Fig. 1F). The intracellular domain of ATRN became critical for Ntx susceptibility in this overexpression assay only when the Ntx dose was lowered near the IC50, suggesting that intracellular trafficking of ATRN is only critical when Ntx concentration is low (fig. S2B). Together, these results indicate that the stem domain of ATRN is critical for its interaction with Ntx.

Knowing the identity of the likely receptor in insect cells, we next investigated whether Ntx could kill mammalian cells expressing the insect ATRN homolog. As previously observed, mammalian cell lines are insensitive to Ntx (*9*). By contrast, HEK293T cells engineered to express GFP-tagged *A. gambiae ATRN* displayed dose-dependent sensitivity to Ntx (Fig. 2A). Moreover, Ntx treatment caused ATRN-GFP to rapidly relocalize to cytoplasmic puncta, consistent with Ntx-triggered endocytosis of ATRN (Fig. 2B). Thus, while mammalian cell lines are not intrinsically sensitive to Ntx, they can be made sensitive by expressing *Anopheles* ATRN. Unexpectedly, expression of the stem domain of the melanocyte mouse ATRN isoform (MmATRN) was sufficient to sensitize Sf9 cells to Ntx, and pulldown assays supported a direct interaction between this MmATRN isoform and Ntx (fig. S2C,D). As ATRN mRNA ubiquitously expressed in mammalian tissues and cell lines, mammalian cell insensitivity to Ntx may be due to a difference in post-transcriptional regulation of ATRN between mammalian and insect cell lines (*13, 18*).

**Fig. 2.**
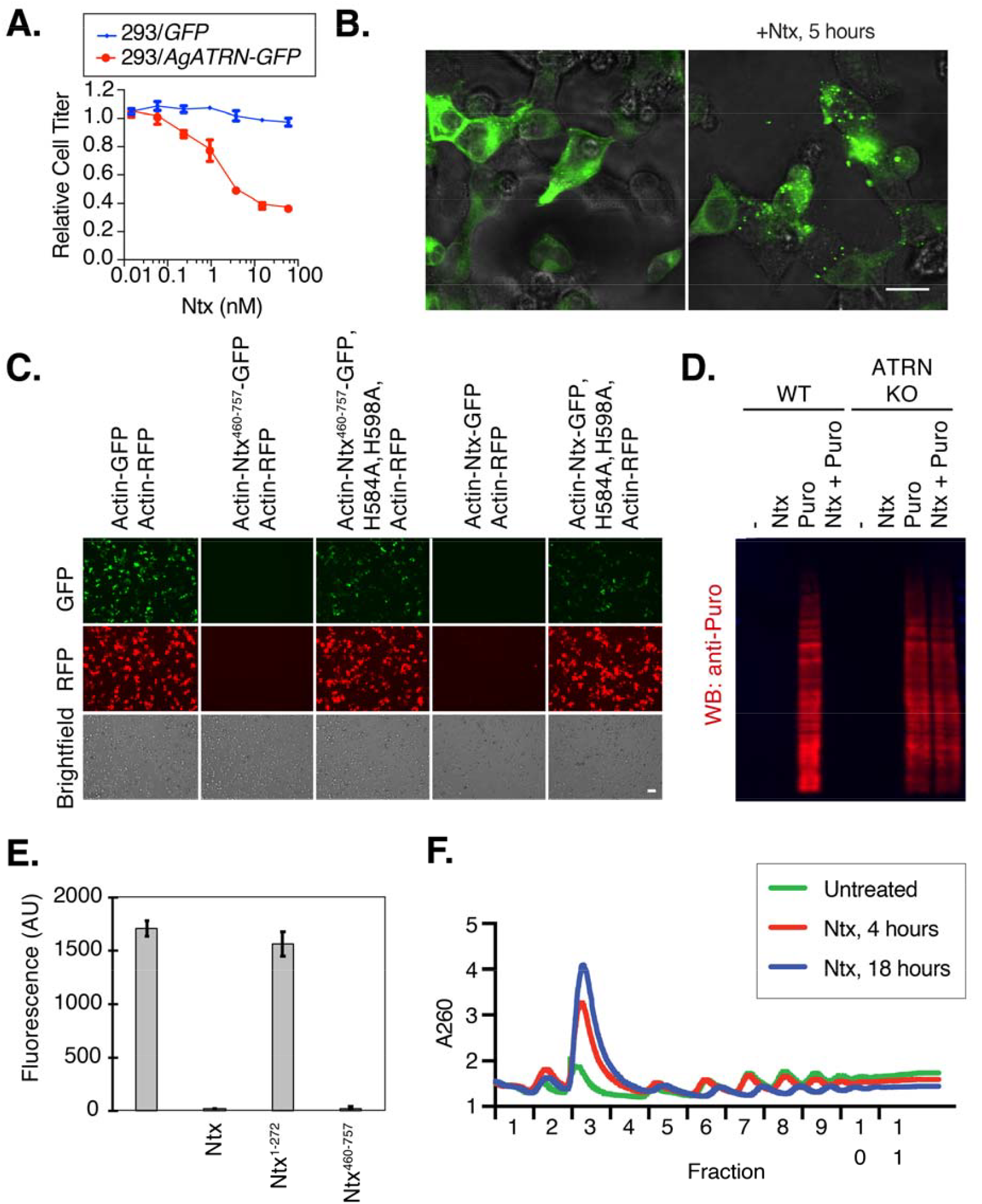
Ntx is an ATRN-dependent inhibitor of protein synthesis. **(A)** Expression of mosquito ATRN sensitizes human HEK293T cells to Ntx. Viability was measured after 48 hours of treatment with increasing doses of Ntx using the Cell Titer Glo assay. Data are means ± SD (*n* = 3). **(B)** Ntx triggers ATRN clustering. Confocal micrographs of HEK293T cells expressing AgATRN-GFP after treatment with 1 nM Ntx for 5 hours. Scale bar = 20 μm. **(C)** The Ntx C-terminal domain inhibits protein expression. Microscope image of cells co-transfected for plasmid expression of the indicated pairs of RFP and GFP tagged proteins. Scale bar = 50 μm. **(D)** Ntx inhibits translation in an ATRN-dependent manner. Wild-type (WT) and *ATRN* KO HEK293T cells were treated with Ntx. Puromycin (Puro) incorporation into nascent proteins was detected by Western blot. **(E)** The Ntx C-terminal domain is sufficient to inhibit translation *in vitro. In vitro* translation reactions in RRL programmed with luciferase mRNA were pretreated with Ntx fragments, and then luminescence was detected. Data are means ± SD (*n* = 3). **(F)** Ntx treatment causes polysome collapse. Ribosome profiles from HEK293T/ATRN-GFP cells treated with 1 nM Ntx for 4 or 18 hours or untreated.

## Ntx inhibits the early steps of protein translation

To explore the downstream effects of Ntx, we determined whether genes in essential processes were partially enriched in the CRISPR screen and noted the presence of genes related to mitochondria and translation (fig. S3A,B). Paradoxically, the downregulation of genes critical for a cellular component serving as the target of a toxin can provide partial resistance to it. For instance, the translation control genes ILF2, ILF3, and RPS25 emerged in an RNAi screen for resistance to the ribotoxin ricin (*19*). As Ntx was previously suspected to act on mitochondria (*9*), we first examined the outcome of Ntx treatment on isolated mitochondria from Sf9 cells using O_2_ consumption assays and found no measured effect (fig. S3C,D). We next examined the impact of Ntx on protein synthesis. In insect cells, transfection with plasmids encoding the GFP-tagged C-terminal domain of Ntx led to the inhibition of its expression as well as that of a co-transfected mCherry reporter gene in the same cells (Fig. 2C). When this C-terminal domain contained mutations H584A and H598A—previously shown to disrupt Ntx killing ability (*9*)— the expression of both mutant Ntx and the reporter was restored (Fig. 2C), supporting a role for Ntx in inhibiting protein synthesis. Moreover, pretreatment of insect cells with Ntx resulted in a failure of puromycin incorporation into nascent peptides, further supporting a primary role in protein synthesis inhibition (Fig. 2D). As a control, *ATRN* knockout cells maintained puromycin incorporation after Ntx treatment, further demonstrating that the ATRN receptor is required for Ntx to access the cytoplasm to inhibit translation (Fig. 2D).

To assess whether Ntx directly affects the translation machinery, we turned to *in vitro* assays. Because we found Ntx activity to be conserved across species, we used rabbit reticulocyte lysate (RRL), a well-established *in vitro* system for studying translation. Incubation of RRL with Ntx led to the elimination of luciferase mRNA translation, with the activity mapping to Ntx’s C-terminal region (Fig. 2E; fig. S4A-C). Consistent with the lack of sequence or structural homology of Ntx with known translation-inhibiting toxins, its inhibition was found to proceed by a unique mechanism, as Ntx was insensitive to nicotinamide treatment (ruling out a diphtheria toxin-like ADP ribosylation function for Ntx) and did not result in depurination of the 28S rRNA (ruling out a ricin-like ribotoxin function) (fig. S4D,E). Moreover, ribosome profiles showed an increase in the 80S monoribosome peak following 4 hours of Ntx treatment, proceeding to a collapse of polyribosomes after 18 hours (Fig. 2F; fig. S4F). This suggests that once in the cytoplasm, Ntx likely inhibits translation initiation, further differentiating its activity from ribotoxins such as ricin, which do not alter ribosome profiles as they prevent all ribosome conformational changes (*20*), or translation elongation inhibitory toxins such as diphtheria toxin, which cause an accumulation of polyribosomes and a decrease of monoribosomes (*21*). These results indicate that once the ATRN-Ntx complex enters the cytoplasm, the Ntx C-terminal domain uses a novel mechanism to inhibit the protein synthesis machinery.

## Targeted mutagenesis in predicted Ntx-ATRN interacting domains abolishes toxicity

To further understand the molecular basis of the ATRN-Ntx interaction, we used AlphaFold Multimer to model the complex. The prediction supported a rigid interaction in which a β-strand within Ntx (residues 221–225) augments a β-sheet in the ATRN stem domain (residues 1015– 1161) (Fig. 3A). β-strand augmentation is a specialized protein-protein interaction mechanism typically employed by signaling proteins such as Notch, Src, and Shc (*22*). Using Predicted Aligned Error (PAE) plots from AlphaFold3, we visualized the loss of interaction by engineering mutations at the β-strand augmentation interface in either Ntx or ATRN (Fig. 3B). Notably, Ntx residues 221-225 are predicted to lose their β-strand formation potential upon mutation of ATRN, supporting the idea that an entropic mechanism drives the interaction.

**Fig. 3.**
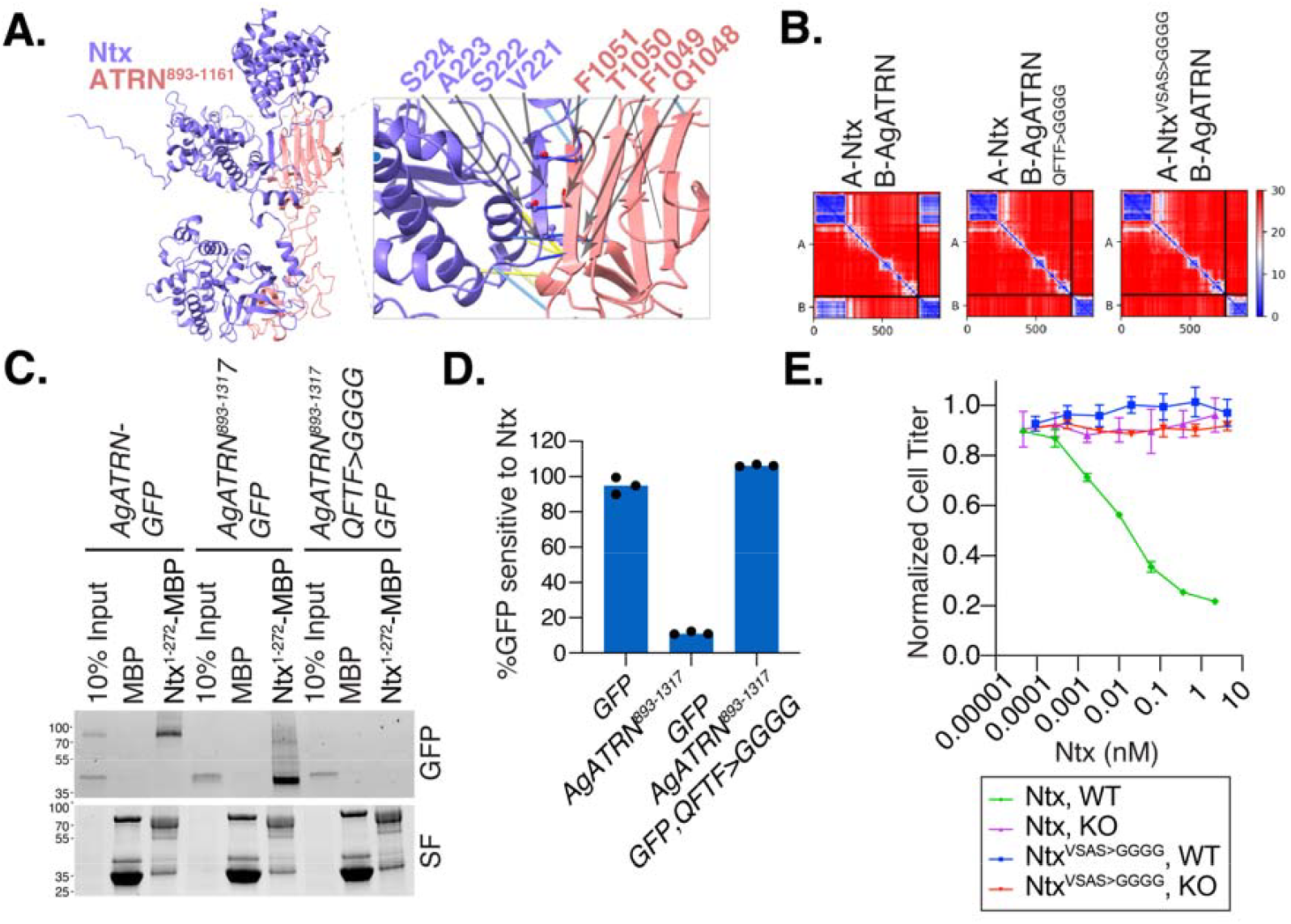
Ntx engages ATRN through β-strand augmentation. **(A)** AlphaFold3 model of the Ntx-ATRN complex. The N-terminal domain of Ntx (purple) interacts with the ATRN stem domain (pink). Inset, close-up of the predicted β-strand augmentation interface highlighting relevant residues. **(B)** Predicted Aligned Error (PAE) plots for wild-type and mutant Ntx-ATRN complexes. Mutations in the interface of either ATRN (QFTF>GGGG) or Ntx (VSAS>GGGG) are predicted to disrupt the interaction. **(C)** Interface mutations abrogate Ntx-ATRN binding. Pulldown of wild-type or mutant (QFTF>GGGG) AgATRN-GFP by the Ntx N-terminal domain (Ntx^1^□^2^□^2^-MBP). SF, stain-free gel. **(D)** Quantification of Ntx sensitivity in Sf9 *ATRN* KO cells expressing wild-type or mutant (QFTF>GGGG) ATRN. Data are means ± SD. **(E)** The Ntx interface is required for cytotoxicity. Viability of WT or *ATRN* KO Sf9 cells treated with increasing doses of wild-type (WT) Ntx or mutant (VSAS>GGGG) Ntx. Data are means ± SD.

Next, we experimentally introduced mutations into the ATRN β-strand augmentation interface and assessed their effect on Ntx binding. As predicted by Alphafold3, immunoprecipitation assays showed complete disruption of the physical interaction between Ntx and the mutant ATRN compared to the wild type (Fig. 3C). In cytotoxicity assays, Sf9 *ATRN* knockout cells expressing mutant ATRN failed to regain sensitivity to Ntx compared to cells expressing wild-type ATRN (Fig. 3D). Conversely, mutating the β-strand augmentation interface in Ntx resulted in the complete loss of its cell-killing activity (Fig. 3E). These findings collectively suggest that β-strand augmentation is a key mechanism mediating the ATRN-Ntx interaction.

## The ATRN binding domain is identified in other Proteobacteria toxins

To determine whether Ntx’s entry domain is shared among other toxins, we performed a search of all public databases for sequence or structural homology to Ntx’s N-terminal domain and modeled the interaction of each candidate with the minimal Ntx-bidning domain, AgATRN(1015-1161), measuring the Alphafold3 interface predicted template modeling (ipTM) score as a prediction of interaction strength with ATRN (Fig. 4A; table S2). The intersection of these approaches identified 60 unique protein groups, which we call <u>A</u>TRN-Interaction <u>D</u>omain-containing <u>P</u>roteins (AIDPs), found exclusively in Gram-negative bacteria, primarily those in the Gammaproteobacteria and a few in Betaproteobacteria, in *Burkholderia* (table S3). Most are identified in the Vibrionaceae and Yersiniaceae, and Enterobacteriaceae. Like Ntx, AIDP genes are typically located in accessory genomic islands and on plasmids (table S4). Neighboring genes contain integrases as well as bacteriophage and T6SS-like contractile secretion system (CIS) (*23*) and *Photorhabdus* virulence cassette (PVC) (*24*) suggestive of horizontal transmission (table S4).

**Fig. 4.**
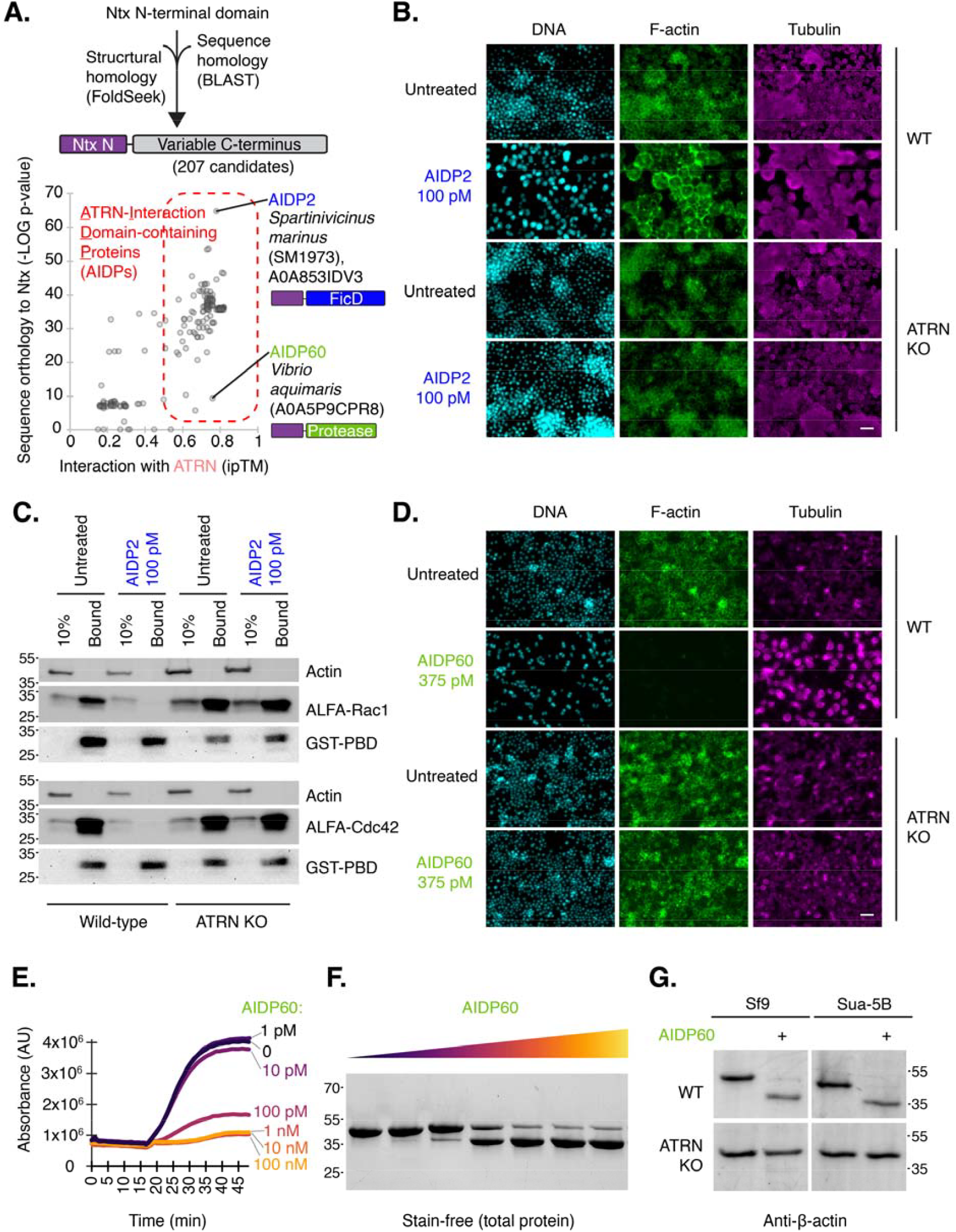
The Ntx entry domain is a scaffold for diverse cytotoxic warheads. **(A)** Identification of <u>A</u>TRN-Interaction <u>D</u>omain-containing <u>P</u>roteins (AIDPs). First, candidates were identified using sequence or structural (FoldSeek) (*40*) homology to Ntx’s N-terminal domain. Next, the interaction with ATRN was predicted by Alphafold modeling, revealing ∼200 AIDPs with variable C-terminal regions, some with homology to known effector domains (table S2 to S5). AIDP2 and AIDP60 are indicated along with the predicted functions of their C-terminal domains. **(B)** AIDP2 alters cell morphology in an ATRN-dependent manner. Representative immunofluorescent staining of WT and *ATRN* KO Sua-5B cells treated with 100 pM AIDP2 or left untreated. DNA is stained with DAPI (cyan), F-actin (green), and tubulin (magenta). Scale bar = 20 μm. **(C)** AIDP2 inactivates Rho GTPases. Lysate from WT or *ATRN* KO Sf9 cells transfected with ALFA-tagged Rac1 or Cdc42 as indicated were subjected to pulldown using GST-tagged p21 binding domain (GST-PBD) to precipitate activated Rho GTPases. ALFA and GST tags were detected by western blot. Antibody against actin serves as a loading control. **(D)** AIDP60 eliminates F-actin in an ATRN-dependent manner. Representative immunofluorescent staining of WT and *ATRN* KO Sua-5B cells treated with 375 pM AIDP60 or left untreated. DNA is stained with DAPI (cyan), F-actin with (green), and tubulin (magenta). Scale bar = 20 μm. **(E)** AIDP60 inhibits actin polymerization *in vitro*. Pyrene-actin fluorescence, reporting actin polymerization state, was measured in the presence of increasing concentrations of AIDP60. **(F)** AIDP60 is an actin protease. Total-protein-stained gel of purified actin was incubated with increasing concentrations of AIDP60. **(G)** AIDP60 cleaves β-actin in cells. Representative anti-β-Actin Western blot of lysates from WT and *ATRN* KO Sf9 or Sua-5B cells treated with AIDP60 or left untreated.

After searching all relevant public databases, no other AIDP had a C-terminal domain with significant similarity to Ntx. Instead, the AIDPs identified had a large diversity of additional domains, typically in the C-terminal region (e.g., transferase, kinase), and some lacked homology with known protein domains (table S5). To ask whether any of these could function as an ATRN-dependent toxin, we first attempted recombinant expression of six candidates in *E. coli*. Two were successfully expressed and purified; we renamed these AIDP2 and AIDP60 (fig. 5A; table S3). Both are predicted to engage ATRN through β-strand augmentation (fig. S5B,C). Treatment with either AIDP led to cell death, and selection of a genome-wide CRISPR guide-expressing population of mosquito Sua-5B cells for resistance to one AIDP resulted cells resistant to the other as well as to Ntx (Fig. 6D,E). Consistent with the observation of cross-resistance, ATRN was the top hit in replicate CRISPR screens with both novel AIDPs (fig. S5F). These data confirm that two AIDPs are novel single-chain cytotoxins that require ATRN to be on host cells to exert their cell killing activities.

We next investigated the mechanism of killing by AIDP2 and AIDP60. AIDP2 from *Spartinivincus marinus* SM1973, a Gram-negative bacterium recently isolated from intertidal sediment (*25*), has a C-terminal Fic domain, which is predicted to catalyze AMPylation and inactivation of Rho GTPases by preventing their binding to effectors (*26*). Fic domains have been found in Gram-negative bacterial toxins that enter cells via Type III, IV, or V secretion systems, but have not been previously described within exotoxins (Type II secreted) (*27*). Alphafold3 modeling shows specific binding of AIDP2 to Rho GTPase Rac1 (fig. S6A). We further confirmed that AIDP2 can transfer AMP to purified Rac1 (fig. S6B). Exposure of cells to AIDP2 resulted in rounding in wild-type but not ATRN-knockout insect cells (Fig. 4B; fig. S6C) and in human cells only if they expressed insect ATRN (fig. S6D). Furthermore, AIDP2 treatment inhibited Rac1 or Cdc42 in wild-type but not ATRN-knockout insect cells (Fig. 4C). Thus, AIDP2 is a toxin possessing the functional entry domain of Ntx together with a Fic AMPylation “warhead”.

AIDP60, from recently described *Vibrio aquimaris* (*28*), also had a profound, ATRN-dependent effect on the cytoskeleton of treated cells; its treatment resulted in the loss of phalloidin staining, indicating near elimination of F-actin with an IC50 of 13 pM ± 27 (Fig. 4D; fig. S7A,B). Consistent with this, addition of AIDP60 to pyrene-actin *in vitro* prevented polymerization of pyrene-actin into F-actin (Fig. 4D). AIDP60’s C-terminal domain has an HExxH metalloprotease motif, so we wondered if it proteolyzes actin directly. Three “actinases,” effectors that cleave monomeric actin, have been previously identified, and actin is a known caspase target; however, actin monomer cleavage has not yet been reported for a receptor-binding exotoxin (*29-31*). Exposing purified actin to AIDP60 *in vitro* led to the detection of a distinctly smaller protein fragment of ∼38 kDa (Fig. 4F). After treatment of cells with AIDP60, we detected an actin form with a similar size-shift. This was observed in wild-type but not ATRN-knockout insect cells (Fig. 4G). Mammalian cells expressing insect ATRN but not parental cells displayed dramatically altered cellular morphology including membrane blebbing, characteristic of cytoskeletal damage, in response to AIDP60 exposure (fig. S7C).

Western blotting with an actin N-terminus antibody showed that cleavage by AIPD60 occurs from the N-but not C-terminus (fig. S7D), similar to protealysin, grimelysin and caspase 3, but unlike RavK (*29-31*) (fig. S7E). The N-terminal 5 residues from the cleavage site were mapped by Edman degradation and identified as a unique site (between Methionine-44 and Valine-45) not shared by any previously characterized actin N-terminus protease (fig. S7F). An actin-cleaving exotoxin would have to be effective at low concentrations, since few molecules of an exotoxin are encountered by host cells and successfully delivered into the cytoplasm. The sub-nanomolar potency of AIDP60 supports this (fig. S7B). We wondered if additional mechanisms exist to restrict AIDP60 activity to actin. Remarkably, Alphafold3 Multimer modeling supports direct binding of AIDP60 to actin at exosite contacts across its exposed surface, positioning the putative catalytic HExxH metalloprotease active site motif at the experimentally identified cleavage site with a high ipTM score, indicative of stable contact (fig. S7G). Binding to exosites far from the cleavage site is a characteristic of two of the most potent exotoxin proteases, botulinum toxin and anthrax lethal factor (*32, 33*). Thus, AIDP60 is a toxin that binds ATRN to gain cell entry and once inside, binds to and site-specifically proteolyzes actin in a manner similar to other highly lethal protease toxins. We hypothesize that this subsequently interferes with a range of host functions including bacterial clearance by immune cells, a key defense against bacterial colonization of the host (*34*).

Here we show that Ntx, a lethal Vibrio exotoxin responsible for devastating shrimp pandemics, uses cell surface protein ATRN to enter host cells via a novel β-strand augmentation interface. Further, we found that Ntx’s C-terminal domain then kills cells by inhibiting translation through a unique mechanism. Ntx may therefore offer a novel scaffold for immunotoxin therapies, in which a lethal toxin is fused with an antibody directing it to target cells (*35*). Moreover, Ntx’s mechanism of translation initiation inhibition may provide added specificity for cancer cells (*36*).

We also found that the ATRN-binding domain of Ntx defines a new family of modular toxins— the AIDPs—in which a conserved entry scaffold is fused to disparate effector domains. The location of AIDP genes on mobile genetic elements likely explains the broad dissemination of this entry scaffold. The functional versatility of this system is exemplified by the distinct enzymatic activities of its members: the translation-inhibiting domain of Ntx; the Rho GTPase-AMPylating Fic domain of AIDP2; and the actin binding and cleaving protease domain of AIDP60. Our findings define a single-chain exotoxin family by its conserved mechanism of cell entry rather than cytotoxic function. Other toxin systems have been found to have multiple effector domains fused to the same polypeptide warhead (*37*), or encode syringe-like nanodevices that allow the entry of diverse effectors through a common pore (*38, 39*), but there has been less evidence that a recognizable receptor binding, translocation domain can be used to identify novel domains associated with effector function. Our work suggests this framework may be generalizable and is likely to have greater value as additional bacterial genome sequences become available.

## Materials and Methods

### Cell Lines and Culture

*Anopheles gambiae* complex (malaria mosquito) Sua-5B cells and *Drosophila melanogaster* S2R+ cells (“S2 cells”) were cultured in Schneider’s Drosophila Medium supplemented with 10% FBS and 1% P/S (Thermo) and maintained in flasks at 25°C. *Spodoptera frugiperda* Sf9 cells were cultured in Sf-900 II SFM supplemented with 10% FBS and 1% P/S (Thermo) and grown in spinner flask culture at 25°C. Human embryonic kidney (HEK) 293T cells were cultured in Dulbecco’s Modified Eagle’s Medium (DMEM; Thermo) supplemented with 10% FBS and 1% P/S and maintained at 37°C in 5% CO□. All cell lines tested negative for mycoplasma contamination.

### Recombinant DNA and Plasmids

Coding sequences for *V. nigripulchritudo* Ntx, AIDP2, and AIDP60 were codon-optimized for *E. coli* expression and synthesized (Genscript). For bacterial expression, toxin genes (full-length Ntx, Ntx(1-272)-MBP, AIDP2, and AIDP60) were subcloned into pET-28a(+) containing an N-terminal His□-SUMO tag. For mammalian and insect cell expression, full-length and fragmented cDNAs for *A. gambiae* ATRN (AgATRN), *L. vannamei* ATRN (LvATRN), mouse ATRN (MmATRN), and Ntx were subcloned into vectors containing baculovirus OpiE2 or Drosophila Actin5C promoter with a C-terminal EGFP tag. *S. frugiperda* ALFA-tagged Rac1 and Cdc42 were synthesized and cloned into pAc5.1. All constructs were verified by Nanopore sequencing (Plasmidsaurus). Complete sequences of all plasmids generated in this study can be found in table S6. Mouse MGRN1-dsRed plasmid was a kind gift of Teresa Gunn (McLaughlin Research Institute).

### Recombinant Protein Expression and Purification

His□-SUMO-tagged recombinant proteins (Ntx, AIDP2, AIDP60) were expressed in *E. coli* BL21(DE3). Cultures were grown at 37°C to an OD□□□ of 0.8–1.0 and induced with 1 mM IPTG for 16 h at 18°C. Cells were lysed by French press at 27 kPa in TBS (20 mM Tris, pH 7.4, 150 mM NaCl, VWR). Proteins were first purified by Ni-NTA affinity chromatography (GE Healthcare). His□-SUMO tag was then cleaved by Ulp1 protease, and untagged proteins were recovered in the flow-through of a second Ni-NTA column. For fluorescence labeling, purified Ntx was labeled using the Mix-n-Stain CF 647 Antibody Labeling Kit (Biotium) and then purified by dialysis in TBS.

### Genome-Wide CRISPR-Cas9 Screen

sgRNAs targeting the *Anopheles gambiae* genome were selected using CRISPR GuideXpress. The top seven sgRNAs per gene were selected based on minimal off-target effect (OTE) score, maximum machine learning (ML) efficiency score, and were filtered to remove sgRNAs matching regions with known SNPs in Sua-5B cell line genome. The library was cloned into a pLib6.4 vector under the control of the Agam_695 U6 promoter (*11, 12*). For library delivery, Sua-5B-IE8-Act::Cas9-2A-Neo cells were transfected with a plasmid mixture containing equimolar amounts of the sgRNA donor plasmid library and an HSP70-ΦC31-Integrase plasmid (pBS130, a gift from Tom Clandin, Addgene #26290) using Effectene (Qiagen) to facilitate recombination-mediated cassette exchange (RMCE). To achieve a coverage of >200 cells/sgRNA, a total of ∼7x10□ cells were transfected. After 4 days, cells were expanded and selected with puromycin. After continuous passaging of the population for 1 month, the cell pool was treated with Ntx (3 nM), AIDP2 (0.5 nM) or AIDP60 (1 nM). Following selection for 2 weeks (Ntx) or 1 month (AIDP2 or AIDP60), genomic DNA was extracted from cell pellets (>1000 cells/sgRNA) using the Quick-gDNA Miniprep kit (Zymo). The genomic DNA was subjected to a 2-step PCR to introduce barcodes and Illumina sequencing adapters. Amplicons were sequenced on a NovaSeq 6000 (Novogene). sgRNAs with low read counts (<10 reads in the plasmid library) were removed from read count files, and enrichment was quantified using MAGeCK (v0.5.9.4). Reported scores are (−1)*Log10(pos(score)) reported by the MAGeCK test subprogram using default parameters.

### Generation of Knockout and Stable Cell Lines

For *Anopheles* Sua-5B cells, stable knockout cell lines were generated by transfecting Cas9-expressing cells with pLib-Agam (puromycin resistance) encoding individual sgRNAs. For *ATRN*, three sgRNAs (5’-CCG GAG GTG TTC GCT CAC TC -3’, 5’-ACG GTG CAG GAG TTT AAC TT -3’, and 5’-AGA TGG GCG GAG CAG TAG TT -3’) were tested; the line generated with 5’-AGA TGG GCG GAG CAG TAG TT -3’ was used for subsequent analysis. For *MGRN1*, three sgRNAs (5’-AGA TCG TGG CAT GCG ACT GG -3’, 5’-CGG TGT CGT CAA GCC GGC TG -3’, and 5’-GAT TCA CTG AGT TAC TGC TT -3’) were tested and found to give similar levels of Ntx resistance; the line generated with 5’-AGA TCG TGG CAT GCG ACT GG -3’ was used for subsequent experiments. Transfected cells were selected in puromycin, and the resulting pools were used directly for experiments. For *S. frugiperda* (Sf9) cells, *ATRN* knockout lines were generated by transfecting cells expressing Cas9 with an sgRNA (5’-TGA GTT CTT CCT GTT CCA AG -3’) targeting the *S. frugiperda ATRN* locus from the Ha3 promoter (*41*). A highly enriched knockout population was obtained by subsequent selection with 1 nM Ntx and validated by re-expression of ATRN, which restored sensitivity. For generating the stable HEK293T cell line, the AgATRN-EGFP coding sequence was cloned downstream of the Tetracycline Response Element 3rd Generation (TRE3G) promoter in the pCW57.1 vector (a gift from David Root, Addgene plasmid # 41393). Lentiviral particles were produced and used to transduce HEK293T cells. Following transduction and selection in puromycin, single-cell clones were expanded, and one clone (2D10) was selected and used for all subsequent experiments. For complementation assays, the pOpiE-2 constructs were transiently transfected into *ATRN* knockout Sf9 cells using TransIT Insect reagent (Mirus).

### Cytotoxicity and Cell Viability Assays

Cells were seeded in 96-well plates (1-3 x 10□ cells/well) and treated with serially diluted toxins for 48 h. Cell viability was measured using the CellTiter-Glo Luminescent Cell Viability Assay (Promega). For competition assays in Sf9 *ATRN* KO cells, viability was assessed by quantifying the percentage of remaining GFP-positive cells using flow cytometry after treatment with 1 nM Ntx.

### Immunoprecipitation and Pulldown Assays

For co-immunoprecipitation, cells were lysed in Pierce IP Lysis buffer (25 mM Tris-HCl (pH 7.4), 150 mM NaCl, 1 mM EDTA, 1% NP-40, and 5% glycerol) supplemented with 1X Protease Inhibitor Cocktail (Sigma). Lysates were incubated with GFP-Trap (Chromotek). For MBP pulldowns, lysates from cells expressing GFP-tagged ATRN constructs were incubated with amylose resin pre-loaded with MBP or MBP-fused Ntx fragments. For active Rho GTPase pulldowns, lysates from cells transfected with ALFA-tagged Rac1 or Cdc42 were incubated with GST-tagged p21 binding domain (PBD) beads (Millipore). Bound proteins were eluted and analyzed by in-gel fluorescence or Western blot.

### Microscopy and Immunofluorescence

Cells seeded on optically clear CellCarrier 384-well plates (Perkin Elmer) were fixed with 4% PFA in Brinkley Buffer 80 (80 mM K-PIPES, pH 6.8, 1 mM MgCl2, 1 mM EGTA), permeabilized with 0.2% Triton X-100, and blocked with 5% BSA. Samples were incubated with primary antibodies, followed by Alexa Fluor-conjugated secondary antibodies and stains (Invitrogen). F-actin was stained with fluorescently-conjugated phalloidin and DNA with DAPI. Tubulin cytoskeleton was stained using mouse-anti-tubulin (T5168, Sigma). Images were captured on an EVOS-M5000 inverted widefield microscope (Thermo) or an In-Cell 6000 (GE) confocal microscope.

### Protein Synthesis Assays

For puromycin incorporation, cells pretreated with or without 1 nM Ntx for 18 hours were incubated with 5 µg/mL puromycin for 2 hours before lysis and processing for Western blot using an anti-puromycin antibody (Millipore, MABE343). All samples were normalized to equal protein amount using the BCA assay. For *in vitro* translation, control protein (maltose binding protein, MBP) or purified toxins were added to nuclease-treated Rabbit Reticulocyte Lysate (Promega, L4960) programmed with firefly luciferase mRNA (ApexBio). Following incubation for the indicated times, luciferase activity was measured to determine translation levels using the Luciferase Assay System (Promega, E1500), and protein levels were analyzed using an antibody (Promega, G7451). For polysome profiling, HEK293T/ATRN-GFP cells were treated with 100 µg/mL cycloheximide, lysed, and layered onto a 10–50% sucrose gradient. Gradients were centrifuged at 39,000 rpm for 2 hours (SW41-Ti rotor) and fractionated with continuous A260 monitoring. Fractions were additionally analyzed by Western blot with rabbit-anti-RpS6 (Cell Signaling Technology, 2217).

### *In Vitro* Activity Assays

For *in vitro* AMPylation, ALFA-Rac1 was obtained by transiently transfecting *Drosophila* S2 cells with pAc5.1/ALFA-Rac using Effectene, followed by ALFA selector purification and elution with ALFA peptide (Nanotag Biotechnologies). The purified ALFA-Rac1 was then incubated with 7 nM AIDP2 and ATP for 1 hour at room temperature, and the reaction was analyzed by Western blot using anti-ALFA and anti-AMP (17G6) antibodies. For actin polymerization assays, ∼10 µM G-actin containing 10% pyrene-labeled actin (rabbit skeletal muscle; Cytoskeleton, Inc.) in G-buffer (5 mM Tris pH 8.0, 0.2 mM CaCl□, 0.2 mM ATP) was pre-incubated with AIDP60 (1 pM to 100 nM) for 1 h at room temperature. Polymerization was then initiated by adding KCl to 50 mM and MgCl□ to 2 mM, and the increase in pyrene fluorescence (Ex: 365 nm, Em: 407 nm) was measured every 60 seconds for 1 hour at 25°C using a SpectraMax luminometer. To visualize actin cleavage, the final reaction products were resolved by SDS-PAGE and visualized with total protein stain (Stain-Free, Bio-Rad). For N-terminal sequencing of the cleavage product, the reaction products were transferred to PVDF membranes, stained with Ponceau S, and the ∼38 kDa cleavage band from treatment with 100 nM AIDP60 was excised and subjected to Edman degradation and Phenylisothiocyanate (PITC) analysis, with products separated by HPLC at the Tufts University Core Facility (TUCF).

### Mitochondrial Respiration

Mitochondria were isolated from Sf9 cells by differential centrifugation. Oxygen consumption was measured using an Oroboros O2k high-resolution respirometer. A substrate-uncoupler-inhibitor titration protocol was used, with sequential additions of pyruvate, malate, ADP, and other reagents to assess different respiratory states.

### Structural Modeling and AIDP searches

Homology searches to identify candidate AIDPs were performed using NCBI BLASTp (cutoff p<0.00001), IMG BLASTp (cutoff p<0.00001), and FoldSeek (*36*), which queries the AlphaFold/Proteome, AlphaFold/Swiss-Prot, AlphaFold/UniProt50, CATH50, GMGCL, MGnify-ESM30, PDB100cutoff databases, e-value<0.05). For BLASTp searches, the sequence of the Nigritoxin N-terminus (1-272) was used for the query. For Foldseek, separate searches were conducted using the Ntx crystal structure (PDB: 5M41) (*9*), truncated to residues 1-267, and the Alphafold3 model of Ntx, 1-272, and the results were combined. A secondary comparative BLAST analysis was then used to remove candidate redundancy based on multiple accession numbers for identical proteins. Other candidate proteins exhibited high sequence similarity to one another. We grouped similar candidates with others if they shared an identity greater than 95%. The protein sequence for each unique candidate was next examined using InterProScan (v5.75-106.0) to identify probable protein families, domains, and motifs, along with their respective approximately defined regions within each protein. ipTM analysis across AIDPs was performed using LocalColabFold v.1.5.5 (*42*), which integrates AlphaFold-Multimer (AFM) v.2.3.1. MSAs were generated using colabfold_search against the UniRef30 (version 2202) and mmseqs2/14-7e284 structural databases. All computations were performed on the Harvard O2 high-performance computing cluster and the Longwood high-performance compute cluster using a variety of NVIDIA GPUs (TeslaV100, RTX6000, RTX8000, A40, A100, L40S, and H100). Each prediction generated five models with five-fold recycling. Structural models of individual protein complexes (Ntx-ATRN, AIDP2/AIDP60-ATRN, AIDP2-Rac1, AIDP60-actin) were generated using AlphaFold3. Alphafold3 Model 0 is shown, although all five models exhibited similar ipTM scores. Structures were visualized using UCSF ChimeraX (version 1.7.1). ipTM and PAE plots from Alphafold were visualized using AFM-LIS (*43*).

## Acknowledgments

We would like to thank Teresa Gunn (McLaughlin Research Institute), and Will Walker for plasmids and helpful discussions; Rakesh Joshi (National Chemical Laboratory, CSIR, India) for helpful discussions; and Grace Kim, Sristhi Goswami, William McKenna, and Kelly Reap for cloning support. We would like to thank Aymelt Itzen (Hamburg University) for the 17G6 anti-AMP monoclonal antibody and Kim Orth and Hilery Gray for facilitating the shipment. Thanks to Matthew Oser and Deli Hong (Dana Farber Cancer Institute) for the anti-actin N-terminus AC-15 antibody. We would like to thank Connie Cepko and Xiang Ma (Harvard Medical School) for HEK293T cells. We are grateful to the Research Computing Group at Harvard Medical School for GPU access in the Harvard O2 high-performance computing cluster and the Longwood high-performance compute cluster. Thanks to the MicRoN (Microscopy Resources on the North Quad) Core for their support & assistance in this work. Edman degradation analysis was conducted at the Tufts University Core Facility. NP is a Howard Hughes Medical Institute (HHMI) investigator.

## Funding

National Institutes of Health grant 5P41GM132087 (NP, SEM)

National Institutes of Health grant 3R01AI170835 (NP)

Quadrangle Fund for Advancing and Seeding Translational Research at Harvard Medical School (NP, JJM)

National Institutes of Health grant R01AI018045 (JJM)

Howard Hughes Medical Institute (NP)

## Author contributions

Conceptualization: RV, WRR, NP, JJM

Methodology: RV, DHL, WRR, ARK, GP, EM, HN

Investigation: RV, DHL, WRR, EM, YH, YH, HN, GP, MB

Visualization: RV, WRR, YH, HN, GP

Funding acquisition: NP, SEM, JJM

Supervision: NP, JJM

Writing – original draft: RV, WRR, NP, JJM

Writing – review & editing: RV, WRR, SEM, DHL, NP, JJM

## Data and materials availability

All data are available in the main text or the supplementary materials.

## Supplementary Materials

Materials and Methods

Figs. S1 to S7

Tables S1 to S6

