## Supplementary_Figures for "A family of lethal exotoxins defined by cell entry via the Attractin receptor"

**Fig. S1.**

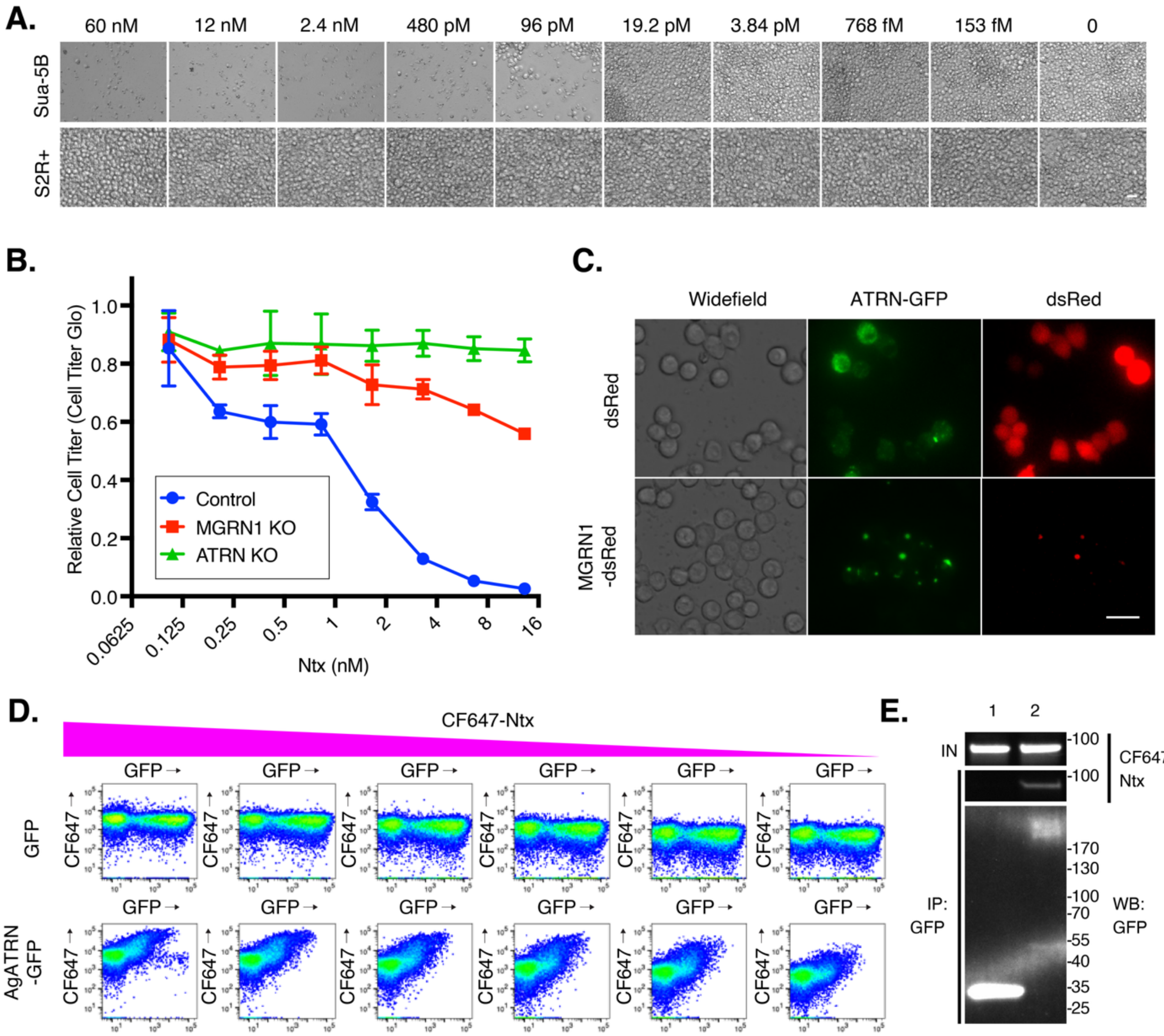


**Characterization of Ntx sensitivity and resistance.** **(A)** Cytotoxicity of Ntx on Sua-5B and S2R+ cells. Brightfield microscopy after 4 days' treatment. Scale bar = 25 µm. **(B)** *MGRN1* KO confers partial Ntx resistance. Cell Titer Glo assay for WT, *MGRN1* KO, and *ATRN* KO Sua-5B cells treated with increasing doses of Ntx. Data are means ± SD. **(C)** MGRN1 alters ATRN localization. Confocal microscopy of S2 cells co-expressing AgATRN-GFP and mouse MGRN1-dsRed. Scale bar = 25 µm. **(D)** Dose-dependent binding of CF647-Ntx to S2 cells expressing AgATRN-GFP, analyzed by flow cytometry. **(E)** Co-immunoprecipitation of Ntx with ATRN. Lysates from cells expressing ATRN-GFP and treated with CF647-Ntx were immunoprecipitated (IP) with anti-GFP antibody. In-gel fluorescence of CF647 and an anti-GFP western blot was performed.

**Fig. S2.**

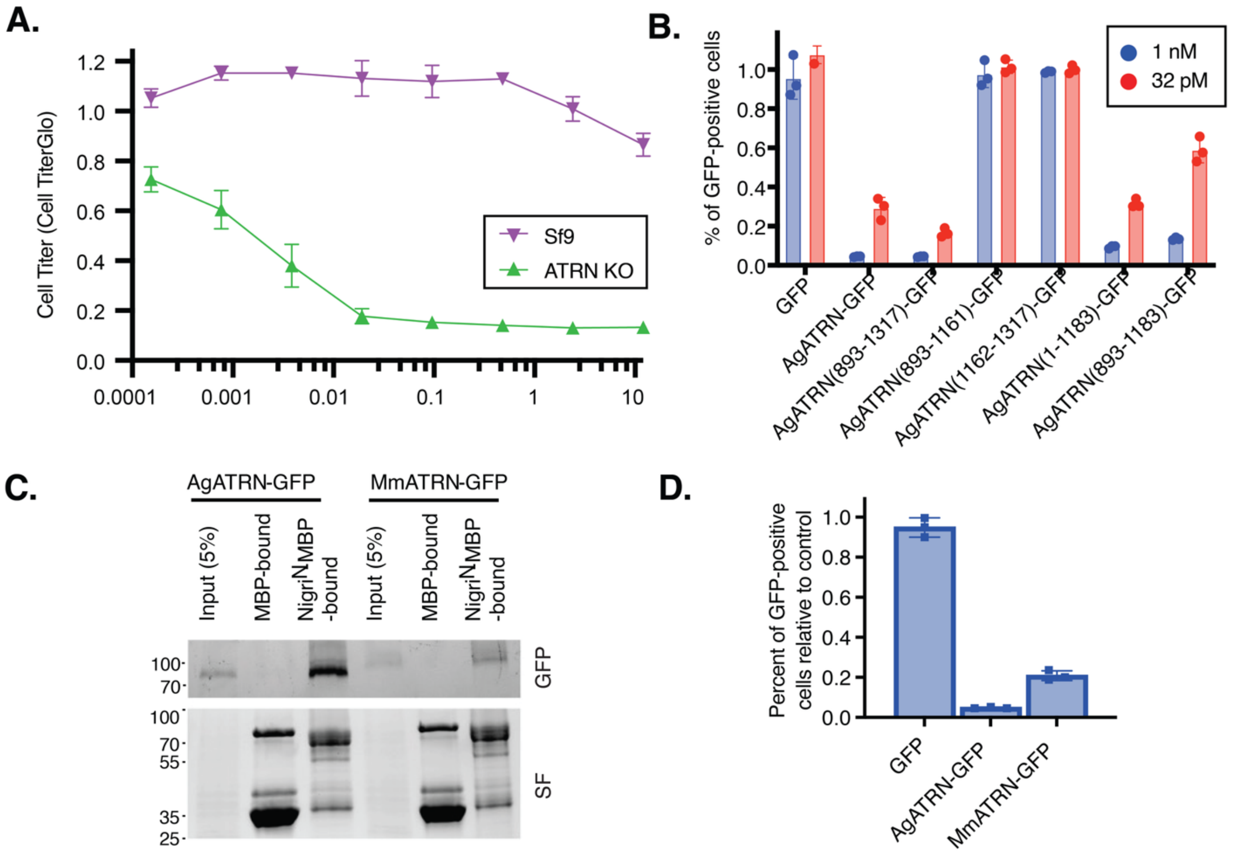


**The ATRN stem domain is a conserved Ntx receptor. (A)** Validation of *ATRN* KO in Sf9 cells. Cell Titer Glo assay for wild-type parental (Sf9) or *ATRN* KO Sf9 cells treated with increasing doses of Ntx. Data are means ± SD. **(B)** Additional mapping of the Ntx-interacting domain of ATRN at 1 nM (blue bars) or 32 pM (red bars) Ntx. Ntx sensitivity of Sf9 *ATRN* KO cells expressing the indicated ATRN constructs. The relative percentage of GFP-positive before vs. after treatmed with 1 nM Ntx was quantified by flow cytometry. Data are means ± SD. Note that ATRN containing a full-length extracellular domain but lacking its intracellular domain (1-1183) or containing a truncated extracellular domain but lacking its intracellular domain (893-1183) fully restores Ntx sensitivity in this overexpression assay when Ntx concentration is high (1 nM, blue bars), but not when it is lowered near the IC50 (32 pM, red bars). **(C)** Ntx binds mouse MmATRN-GFP. Pulldown of AgATRN-GFP or MmATRN-GFP expressed in Sf9 cells by the N-terminal domain of Ntx (Ntx1-272-MBP). **(D)** The mouse ATRN ectodomain partially restores Ntx sensitivity to Sf9 *ATRN* KO cells. The percent of GFP-positive cells following 1 nM Ntx relative to untreated cells (control) measured by flow cytometry is plotted. Data are means ± SD.

**Fig. S3.**

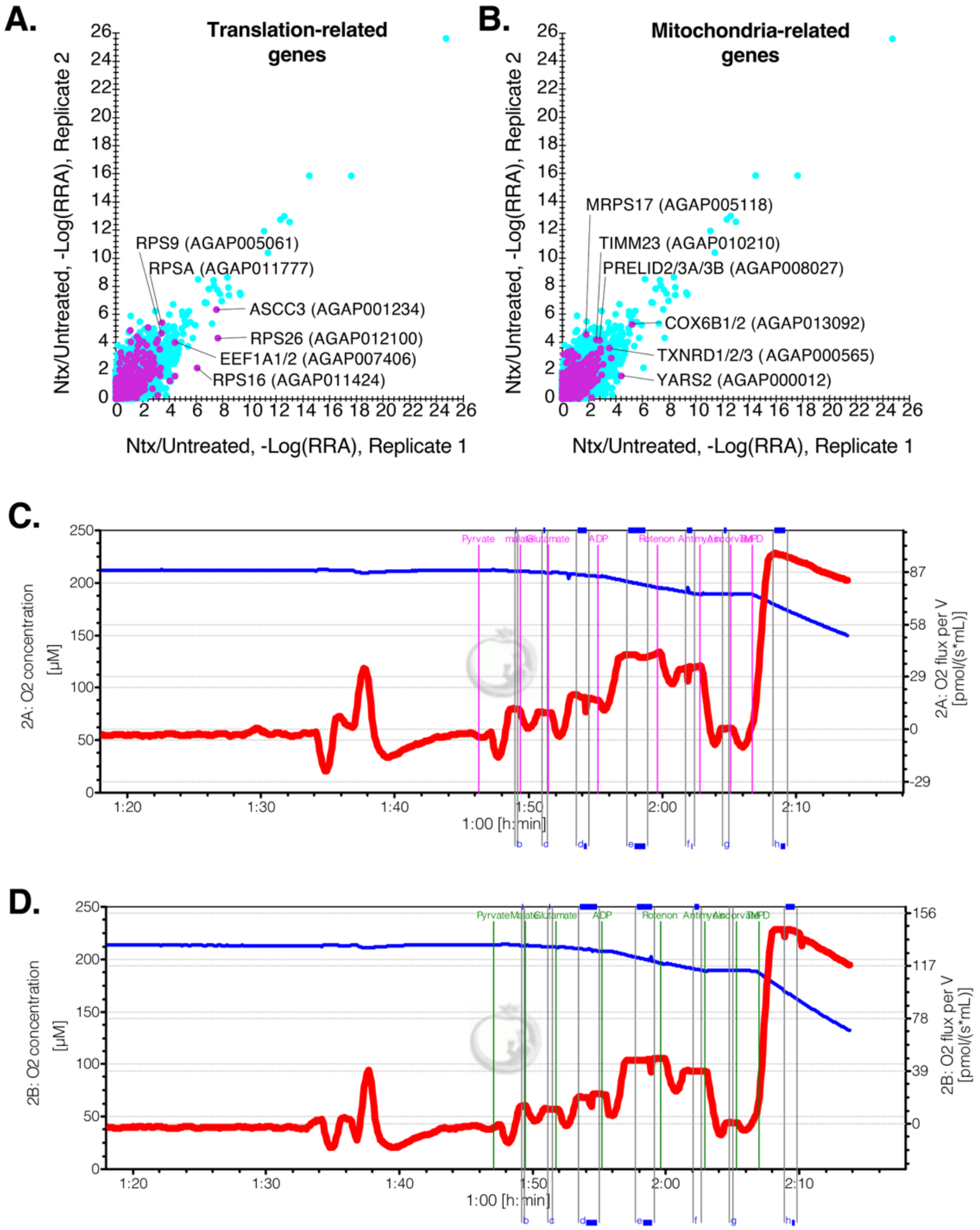


**Minor screen hits include genes involved in translation and mitochondrial functions. (A and B)** CRISPR screen hits related to translation (A) and mitochondria (B). All hits are blue dots. Genes from each indicated category are purple dots. **(C and D)** Ntx does not affect mitochondrial respiration. High-resolution respirometry (Oroboros O2k) traces from isolated Sf9 mitochondria from untreated (C) and Ntx-treated (D) cells. The blue line represents $O_2$ concentration (left y-axis, [µM]), and the red line represents real-time $O_2$ flux (right y-axis, [pmol/s*mL]). Vertical lines indicate sequential addition of mitochondrial substrates and inhibitors to assess different respiratory states.

**Fig. S4.**

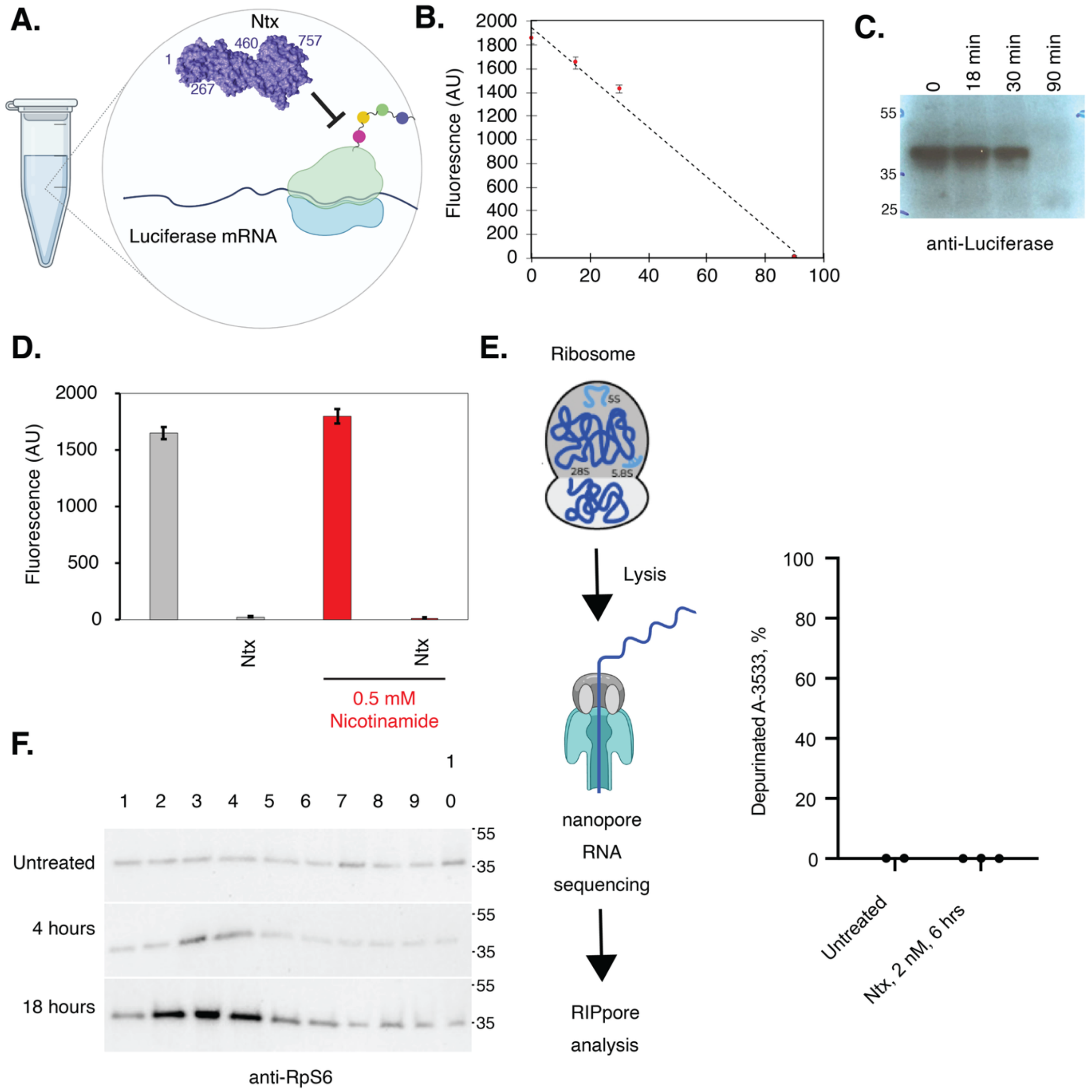


**Ntx inhibits translation via a novel mechanism. (A)** Schematic of the *in vitro* translation assay. Ntx (purple) is shown inhibiting a ribosome translating luciferase mRNA (black squiggle) in a rabbit reticulocyte lysate (RRL) system. **(B)** Time course of *in vitro* translation inhibition by Ntx. **(C)** Western blot of luciferase protein from the in vitro translation assay. **(D)** Ntx activity is not dependent on ADP-ribosylation. *In vitro* translation assay showing that the addition of 0.5 mM nicotinamide, an inhibitor of ADP-ribosyltransferase, does not rescue Ntx-mediated inhibition of luciferase synthesis. **(E)** Nanopore RIPpore analysis (*35*), directly measuring the level of depurination of 28S rRNA directly shows no significant elevation in 28S rRNA depurination after a 6 hour treatment of Sf9 cells with Ntx. **(F)** Ntx treatment causes an increase in monosome fraction at the expense of polysomes, as visualized by the presence of the small ribosomal subunit in the fractions. Western blot for small ribosomal subunit component RpS6 in fractions from a sucrose gradient of Ntx-treated cells.

Fig. S5.

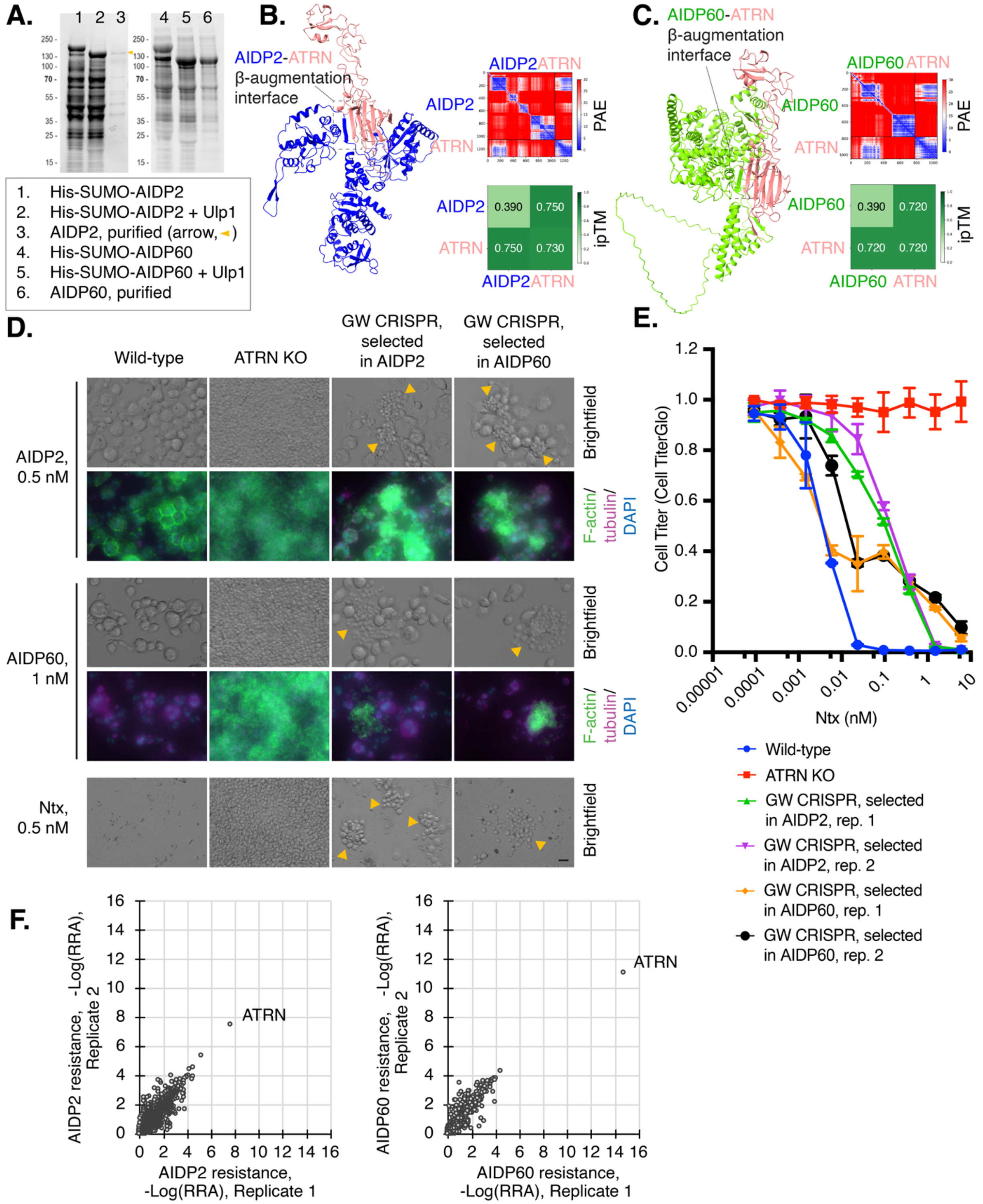


**AIDP2 and AIDP60 are ATRN-dependent toxins. (A)** Purification of recombinant AIDP2 and AIDP60. Total protein stained SDS-PAGE gel of purified proteins before and after $His_6$-SUMO tag cleavage. **(B and C)** AlphaFold3 models of AIDP2-ATRN (B) and AIDP60-ATRN (C)

complexes. PAE and ipTM plots are shown and indicate a rigid interaction. Box: Ntx-ATRN β-augmentation interface. **(D)** Selection of toxin-resistant cell populations. Brightfield and fluorescence micrographs of wild-type, *ATRN* KO Sua-5B, or a genome-wide CRISPR library in Sua-5B cells after selection with AIDP2 or AIDP60, and re-treatment with Ntx, AIDP2, or AIDP60. AIDP2 and AIDP60 induce large, multinucleate cells. Resistant colonies emerge from the selected populations and in *ATRN* KO cells that maintain normal morphology (yellow arrowheads). Scale bar = 20 µm. **(E)** Cross-resistance of selected cell populations. Dose-response curves for cells selected for resistance to AIDP2 or AIDP60 and retreated with Ntx at varying doses. **(F)** CRISPR guide sequencing and analysis from cells selected for 2 months in 0.5 nM AIDP2 or 1 nM AIDP60 reveals that ATRN is the top hit from a genome-wide CRISPR screen for AIDP2 or AIDP60 resistance.

**Fig. S6.**

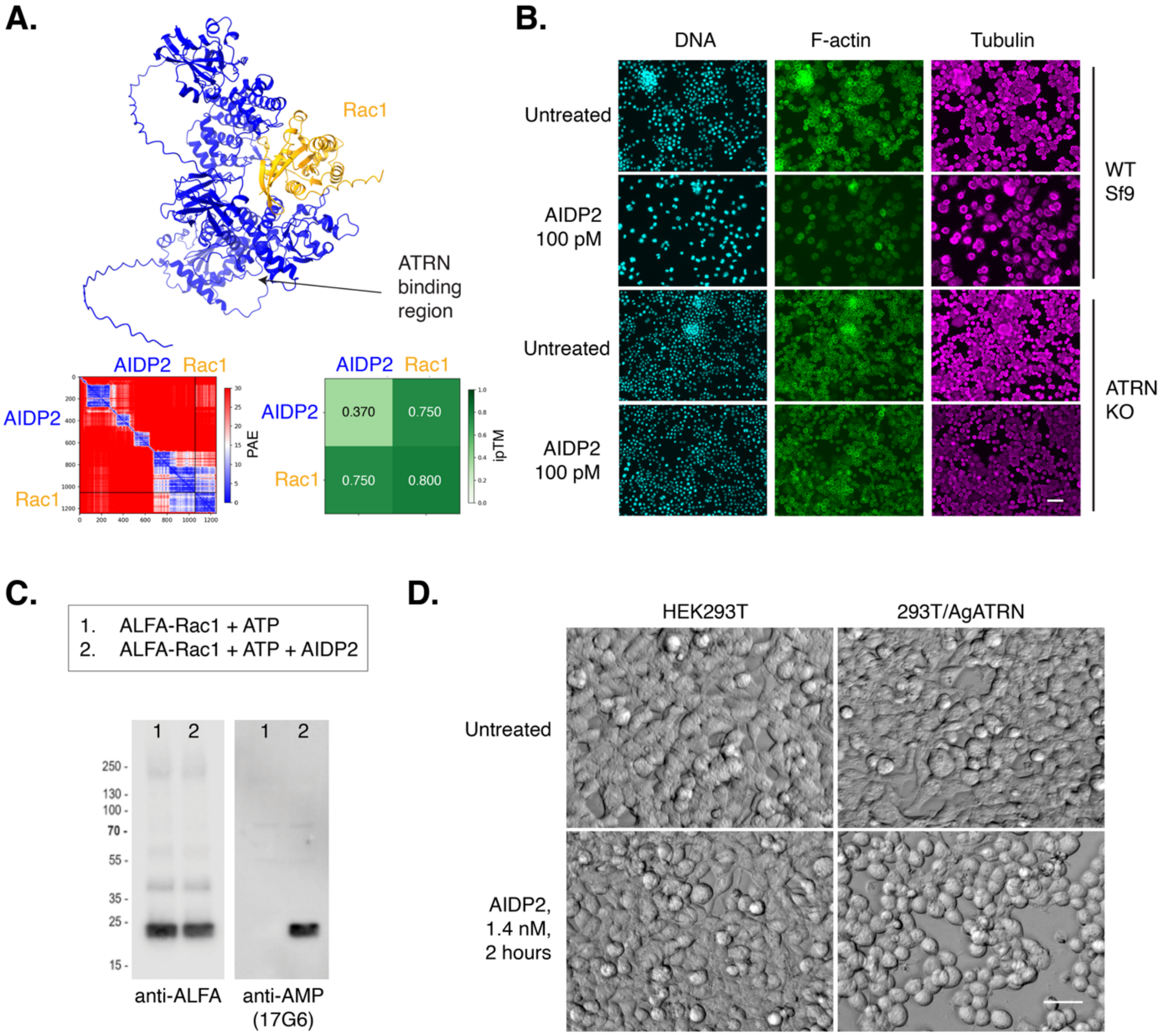


**AIDP2 is an ATRN-dependent Rho-AMPylating toxin. (A)** AlphaFold3 model of the AIDP2-Rac1 complex showing extensive binding of AIDP2 (blue) to Rac1 (yellow). PAE and ipTM plots are shown and indicate a rigid interaction. **(B)** AIDP2 AMPylates Rac1 *in vitro*. AIDP2 (7 nM) was added to purified ALFA-Rac1 and subjected to Western blot using an anti-ALFA antibody or the anti-AMP monoclonal antibody 17G6. Scale bar = 20 µm. **(C)** Cytoskeletal disruption by AIDP2 in WT and *ATRN* KO Sf9 cells. Immunofluorescence of WT and *ATRN* KO Sf9 cells treated with 100 pM AIDP2. DNA is stained with DAPI (cyan), F-actin (green), and tubulin (magenta). **(D)** AIDP2 cytotoxicity requires ATRN expression in mammalian cells. Parental HEK293T or HEK293T/AgATRN-GFP cells were subjected to AIDP2 (1.4 nM) for 2 hours and imaged by brightfield, revealing rounded morphology after treatment with AIDP2. Scale bar = 20 µm.

Fig. S7

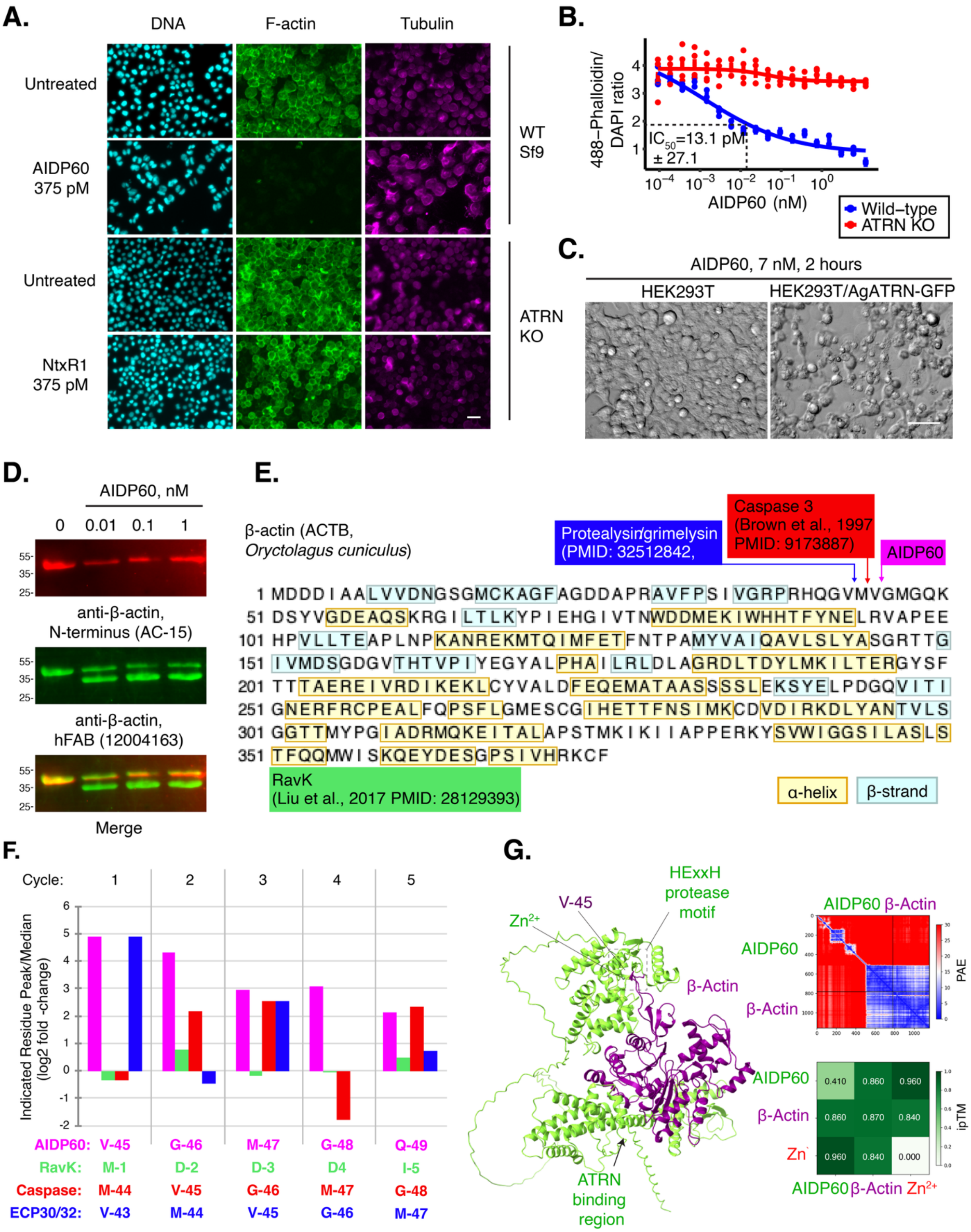


**AIDP60 is an actin-cleaving protease exotoxin. (A)** Immunofluorescence of WT and *ATRN* KO Sf9 cells treated with AIDP60. DNA is stained with DAPI (cyan), F-actin (green), and tubulin (magenta). Scale bar = 20 μm. **(B)** Quantification of phalloidin intensity normalized to DAPI intensity from WT and *ATRN* KO treated with varying concentrations of AIDP60. The data were fitted to a four-parameter logistic (variable-slope) nonlinear regression of the response vs. log[concentration] curve to identify the IC50. **(C)** AIDP60 cytotoxicity requires ATRN expression in mammalian cells. Parental HEK293T or HEK293T/AgATRN-GFP cells were subjected to AIDP60 (7 nM) for 2 hours and imaged by brightfield. While there were no obvious changes in the parental cells, AgATRN-GFP cells exhibited dramatically altered morphology, including membrane blebbing. Scale bar = 20 μm. **(D)** AIDP60 cleaves the N-terminus of β-actin. Western blots of rabbit (*Oryctolagus cuniculis*) β-actin treated with AIDP60 at indicated time for 2 hours at room temperature. Western blot with AC-15 (red) or unmapped monoclonal hFAB (green) anti-β-actin antibodies are shown. **(E)** Mapping the AIDP60 cleavage site on rabbit β-actin. Cleavage site of known actin-cleaving proteases are indicated. Yellow residues = alpha helix; blue = beta sheet. **(F)** HPCL analysis of Edman N-degradation of purified larger actin cleavage product reveals the site V-G-M-G-Q, distinguishing it from other actin cleaving proteases. **(G)** AlphaFold3 model of the AIDP60-β-actin complex revealing extensive contact between the exposed surface of β-actin and the C-terminal region of AIDP60. Putative protease domain is positioned between M-44 and V-45, the experimentally defined cleavage site. PAE and ipTM plots are shown and indicate a rigid interaction.

### Supplementary Tables Legend

**Table S1. (separate file)**
Ntx CRISPR screen results. sgRNA level or Gene-level results of whole-genome CRISPR screens from *Anopheles* Sua-5B cells.

**Table S2. (separate file)**
Results of sequence and structural homology search to Ntx N-terminal domain across all available sequence databases. Candidate protein accession numbers and the corresponding bacterial genera and species. Similar candidates are grouped when protein identity was greater than 95%. For ipTM > 0.50, the protein is listed as an AIDP (ATRN-interaction domain-containing protein). The approximate size and predicted domains of each C-terminal domain (CTD) are provided. BLASTp scores (presented as -Log(p-value)) and Alphafold ipTM scores are listed.

**Table S3. (separate file)**
Collapsed representatives for each unique AIDP. Gray AIDPs = failed purification as SUMO fusions in E. coli. Yellow AIDPs = successfully purified as SUMO-fusions in E. coli.

**Table S4. (separate file)**
Genes located up to 5kb upstream and downstream of each AIDP candidate protein gene are provided. For genome accession contigs that fail to include 5kb on either side, all genes present are identified.

**Table S5. (separate file)**
Table showing all motifs, domains, and families identified and their location in AIDP candidate proteins using InterproScan.

**Table S6. (separate file)**
Sequences of new plasmids generated for this study.
